## Supplementary Information, tables for "Topological signatures in regulatory network enable phenotypic heterogeneity in small cell lung cancer": SI_Oct 30_final.pdf

### Supplementary Materials and Methods

Boolean framework – Ising model with Asynchronous update:

Ising model formalism uses discrete variables to represent the expression level of molecular species (such as micro-RNA or transcription factors etc.). Therefore, the state of the regulatory network (N-node network) can be represented by a sequence {$s_i$, $s_i \in$ {-1, 1}} called a "Boolean Vector" of N binary variables where $s_i$ = 1 represents high-expression level of *i* th node and $s_i$ = -1 represents low-expression of the node. In modeling the dynamics of network via this framework, the only knowledge required is whether each regulatory relationship between network nodes is activating or inhibitory. Regulatory interactions between the molecular species are represented by an N*N matrix called an 'Interaction Matrix (M)' where $M_{ij}$ = 1 represents 'promotion' (or activation) of levels of *i*-th node by *j*-th node and $M_{ij}$ = -1 represents 'inhibition' (or repression) of levels of *i*-th node by *j*-th node of the N-node network. The absence of any regulatory relationship between species *i* and species *j* is indicated by $M_{ij}$ = 0.

At every discrete time step, the expression level of a node $s_i$(t+1) is given as +1 if $\sum_{i=1}^{N} M_{ij} * s_j(t) > 0$; -1 if $\sum_{j=1}^{N} M_{ij} * s_j(t) < 0$; and remains the same when $\sum_{j=1}^{N} M_{ij} * s_j(t) = 0$. The expression levels are updated using the Asynchronous scheme in which a node from the N-node network is picked up at random at every discrete time step and updated using the above non-linear relation. For large discrete-time dynamics, the network settles in a steady state which means that the Boolean Vector is a fixed point of the above-given relation (i.e. will not change as time progresses).

RAndom CIrcuit PErturbation (RACIPE):

RACIPE is a tool which identifies robust dynamical properties of transcriptional regulatory networks (TRNs) by generating an ensemble of continuous network models with distinct kinetic parameters. For every continuous model of a TRN, RACIPE first generates a system of Ordinary Differential Equations (ODEs). For a given node 'N' of the TRN and a set of input activating edges $A_i$ and input inhibiting edges $I_j$, the differential equation corresponding to the expression level of N is given as:

$$\frac{\mathrm{d}N}{\mathrm{d}t} = G_N * \prod_i \frac{H^{S+}(A_i, A^0_{iN}, n_{A_i,N}, \lambda_{A_i,N})}{\lambda_{A_i,T}} * \prod_j H^{S-}(I_j, I^0_{iN}, n_{I_j,N}, \lambda_{I_j,N}) - k_N * N$$

Here, $N, A_i$ and $I_j$ represent the expression levels of the species of the TRN. $G_N$ and $k_N$ denote the Production and Degradation rates respectively. ${A^0}_{iN}$ is the threshold value of $A_i$ expression level at which the non-linearity in the dynamics of $N$ due to $A_i$ is seen. $n$ is termed as the Hill-coefficient and represents the extent of non-linearity in the regulation. $\lambda$ represents the fold change in the target node expression level upon over-expression of regulating nodes. The functions $H^{S+}$ and $H^{S-}$ are known as Shifted Hill functions (Lu *et al.* PNAS 2013) and represent the regulation of the target node by the regulatory node. Shifted hill functions take the following form:

$$H^{S\pm}(B, B^0_A, n_{B,A}, \lambda_{B,A}) = \frac{B_A^{0n_{B,A}}}{B_A^{0n_{B,A}} + B^{n_{B,A}}} + \lambda * \frac{B^{n_{B,A}}}{B_A^{0n_{B,A}} + B^{n_{B,A}}}$$

For the system of ODEs, RACIPE randomly samples the kinetic parameters from a pre-defined set of parameter ranges. At each parameter set, RACIPE integrates the model from multiple initial conditions and obtains steady states in the state space. For the current analysis, a sample size of 10^6 for parameters sets and 1000 for initial conditions was used. The parameters were sampled via a uniform distribution and the ODE integration was carried out using the Runge-Kutta-Fehlberg method of numerical integration.

For the given TRN with ‘n’ nodes, the steady-state expression levels of the nodes were normalized in the following way:

$$E_{in} = \frac{E_i}{f_i}$$

$$f_i = \frac{g_i}{k_i} \prod_j \lambda_{ij}$$

For the i-th node, $E_{in}$ is the normalized expression level of the node, $E_i$ is the steady-state expression level, $f_i$ is the normalization factor, $g_i$ and $k_i$ are the production and degradation of the i-th node corresponding the current steady-state and $\lambda_{ij}$ are the fold change in the expression of node i due to node j. The normalized expression levels of all steady states are then converted into z-scores by scaling about their combined mean:

$$Z_i = \frac{E_i n - \overline{E_{in}}}{\sigma_{in}}$$

where $\underline{E_{in}}$ is the combined mean and $\sigma_{in}$ is the combined variance.

The z-scores can then be classified into Low (zero) and High (one) expression levels based on the sign of their values. This way we have discretized the continuous steady-state levels of the network for comparison with the frequency of Boolean steady states. The way to calculate the total frequency of each discrete state is by counting the occurrence in all the parameter sets. For parameter sets with n steady states, the count of each steady state is taken as 1/n, invoking the assumption that all the states are equally stable.

### Supplementary Figures

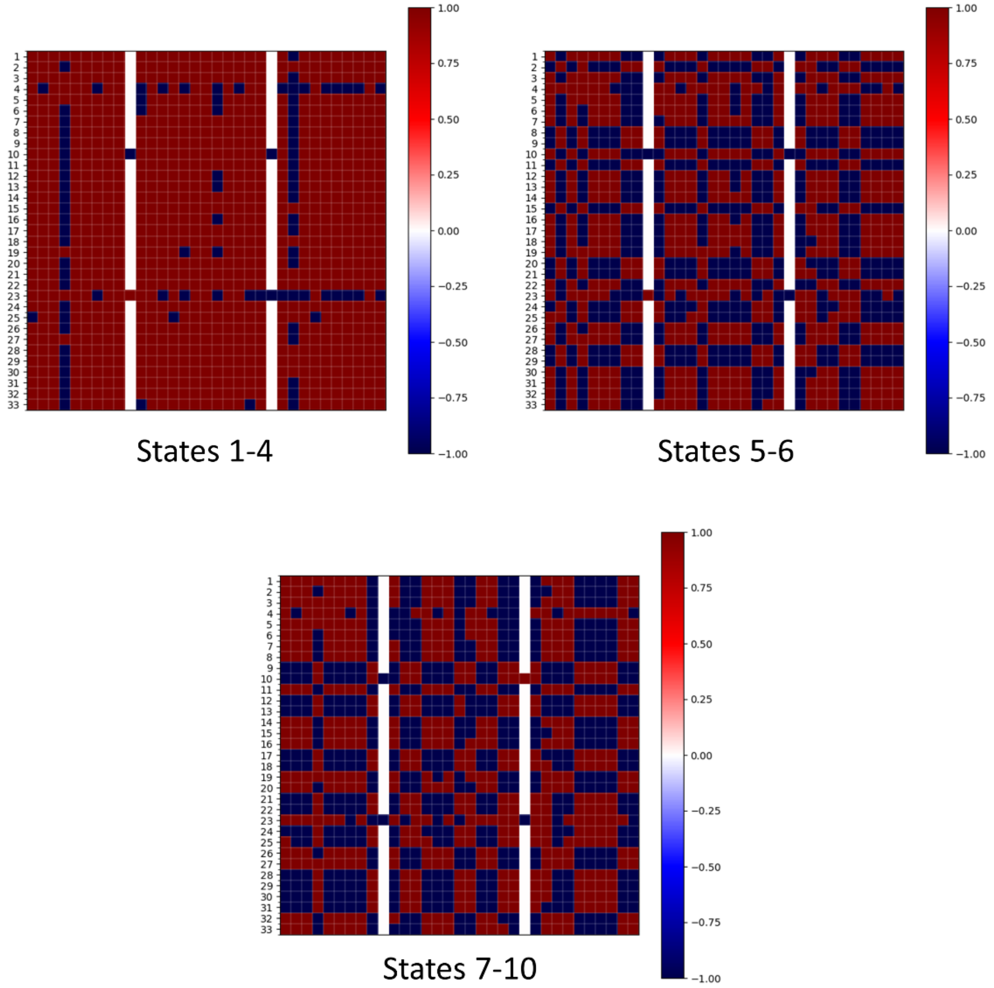


***Fig S1: Influence matrix based on Boolean simulations for states with various frustration levels**. Influence matrix in Fig 3A, i is based on simulations from RACIPE. Influence matrices based on Boolean simulation results were also derived. Instead of taking all steady-state solutions together, we classified them into 3 categories – States 1-4 in Table S1 (frustration value =0.142), states 5-6 (frustration = 0.375) and states 7-10 (frustration = 0.383-0.386) – and derived corresponding influence matrices shown above. Labels 1-33 are the same as in Fig 2. The larger the frustration (or the smaller the frequency), the more the deviation of influence matrix from the correlation matrix shown in Fig 2.*

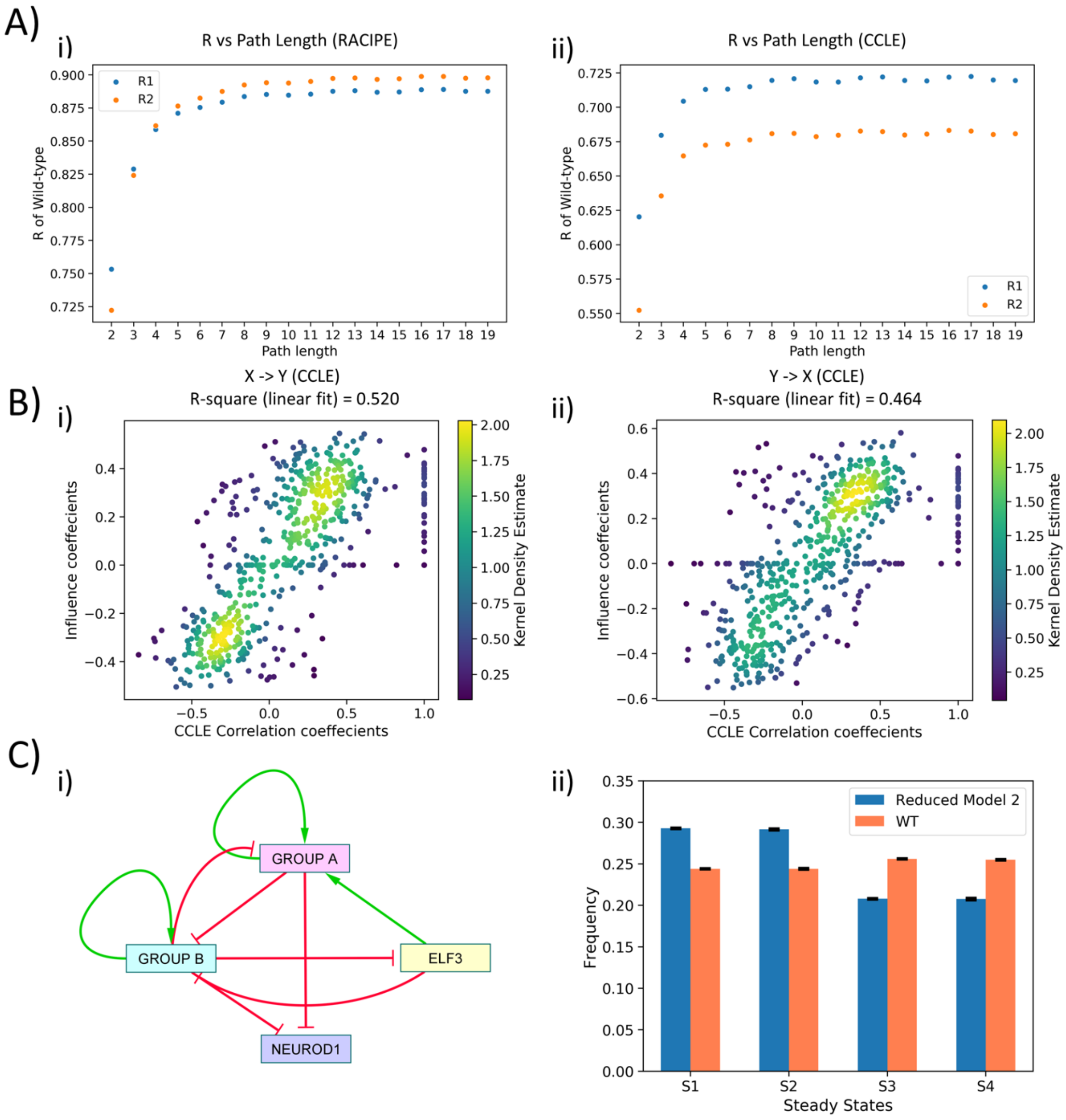


***Fig S2: Topological features**. **A)*** *Scatter plot of R1 and R2 values obtained for varying path lengths, when influence matrix values/coefficients (for varying path lengths) for WT SCLC network were regressed against correlation coefficients from i) RACIPE, and ii) CCLE.* ***B)*** *i) Scatter Plot of R1 and R2 values of influence coefficients of WT network and CCLE correlation coefficients for path length = 10.* ***C)*** *i) Reduced model derived from interaction matrix; red bars show inhibition, green arrows show activation. ii) Steady-state frequencies obtained from reduced model 1 and that of WT SCLC network (n=3).*

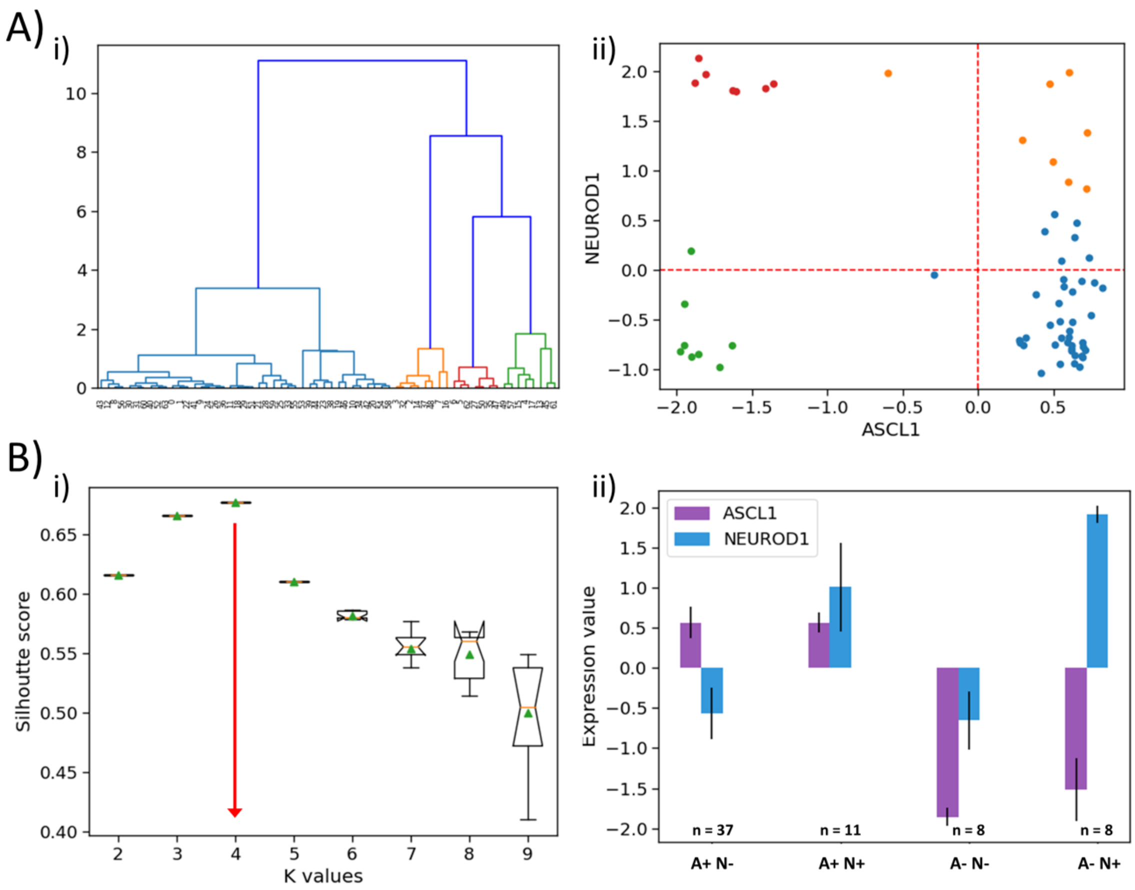


***Fig S3: Analysis for GSE73160 using ASCL1 and NEUROD1. A)*** *i) Hierarchical clustering of GSE73160 dataset across ASCL1 and NEUROD1). ii) Scatter plot of normalized gene expression for NEUROD1 vs ASCL1 of GSE73160 dataset, labeled by n=4 clusters obtained from the earlier dendrogram.* ***B)*** *i) Silhouette score analysis across ASCL1 and NEUROD1 for the GSE73160 dataset. ii) Expression levels ASCL1 and NEUROD1 in 4 clusters obtained from the GSE73160 dataset by k-means clustering.*

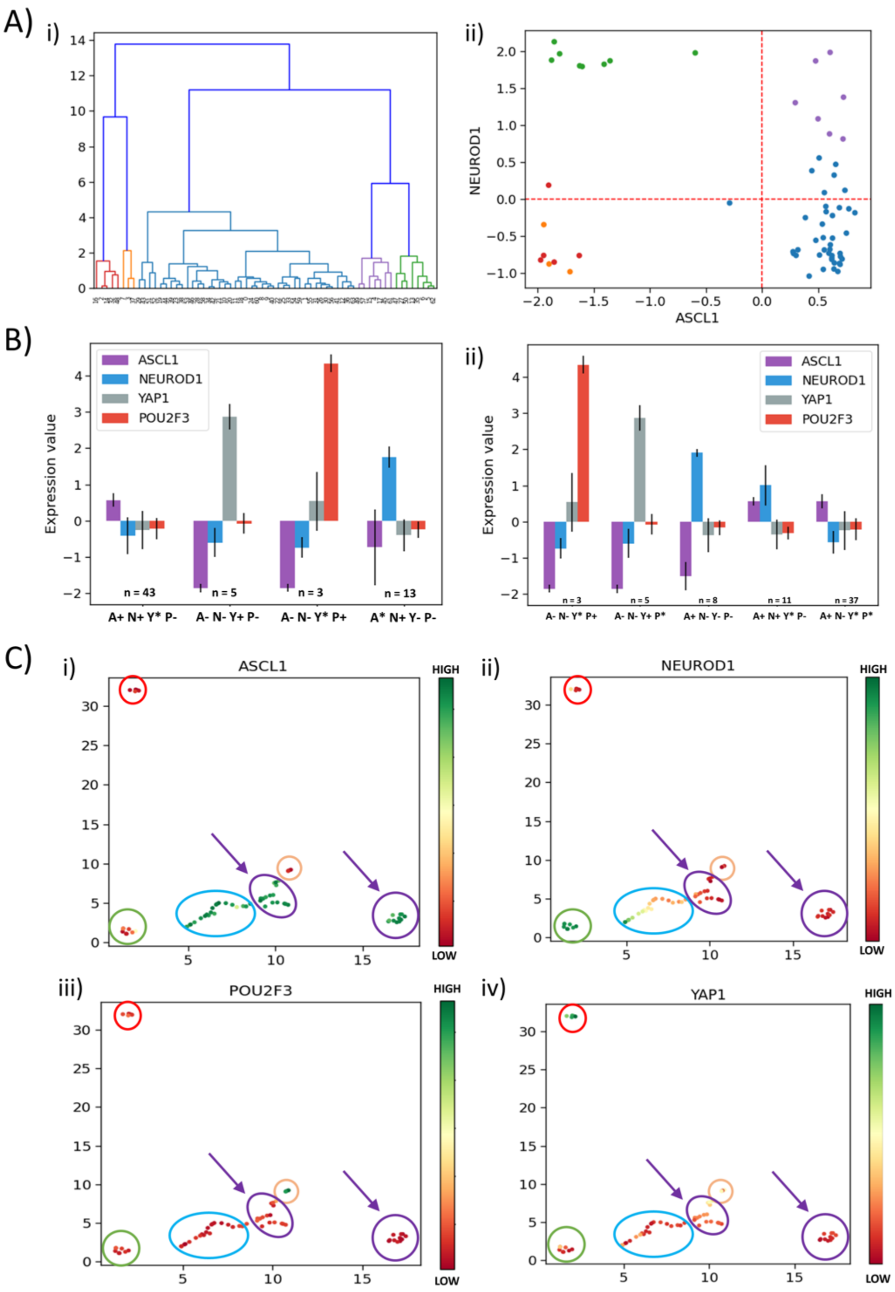


***Fig S4: Analysis for GSE73160 using ASCL1, NEUROD1, POU2F3 and YAP1. A)*** *i) Hierarchical clustering of GSE73160 dataset across ASCL1, NEUROD1, YAP1 and NPOU2F3). ii) Scatter plot of normalized gene expression for NEUROD1 vs ASCL1 of GSE73160 dataset, labeled by n=5 clusters obtained from the earlier dendrogram.* ***B)*** i) Expression levels of 4 classifying genes (ASCL1, NEUROD1, YAP1, and POU2F3) in clusters obtained from the k-means algorithm for k=4 in the GSE73160 dataset. ii) Same as i) but for k=5. **C)** i)-iv) UMAP projection of GSE73160 dataset from 4 dimensions (ASCL1, NEUROD1, YAP1 and POU2F3) to two dimensions. The labeling is based on expression levels of ASCL1 (i), NEUROD1 (ii), POU2F3 (iii), and YAP1 (iv).
